# Control of synaptic transmission and neuronal excitability in the parabrachial nucleus

**DOI:** 10.1101/2020.10.01.322131

**Authors:** Nathan Cramer, Gleice Kelli Silva-Cardoso, Asaf Keller

**Affiliations:** Department of Anatomy and Neurobiology and the Program in Neuroscience, University of Maryland School of Medicine, Baltimore, MD, 21201

**Author notes:** Corresponding Author: Nathan Cramer, Ph.D., University of Maryland School of Medicine, Baltimore, MD 21201. Department of Psychology, Faculty of Philosophy, Sciences and Letters of Ribeirão Preto, University of São Paulo, Brazil (Gleice Kelli Silva-Cardoso).

## Abstract

The parabrachial nucleus (PB) is a hub for aversive behaviors, including those related to pain. We have shown that the expression of chronic pain is causally related to amplified activity of PB neurons, and to changes in synaptic inhibition of these neurons. These findings indicate that regulation of synaptic activity in PB may modulate pain perception and be involved in the pathophysiology of chronic pain. Here, we identify the roles in PB of signaling pathways that modulate synaptic functions. In pharmacologically isolated lateral PB neurons in acute mouse slices, we find that baclofen, a GABA_B_ receptor agonist, suppresses the frequency of miniature inhibitory and excitatory postsynaptic currents (mIPSCs and mEPSC). Activation of µ-opioid peptide receptors with DAMGO had similar effects, while the k-opioid peptide receptor agonist U-69593 suppressed mIPSC release but had no consistent effects on mEPSCs. Activation of cannabinoid type 1 receptors with WIN 55,212-2 reduced the frequency of both inhibitory and excitatory synaptic events, while the CB1 antagonist AM251 had opposite effects on mIPSC and mEPSC frequencies. AM251 increased the frequency of inhibitory events but led to a reduction in excitatory events through a GABA_B_ mediated mechanism. Although none of the treatments produced a consistent effect on mIPSC or mEPSC amplitudes, baclofen and DAMGO both reliably activated a postsynaptic conductance. Together, these results demonstrate that signaling pathways known to modulate nociception, alter synaptic transmission and neuronal excitability in the lateral parabrachial nucleus and provide a basis for investigating the contributions of these systems to the development and maintenance of chronic pain.

**Highlights:**

- *The parabrachial nucleus (PB) is a hub for processing interoceptive and exteroceptive noxious stimuli, including pain*.
- *Synaptic activity in PB is abnormal in chronic pain*.
- *Synaptic activity in PB is regulated by presynaptic and postsynaptic* GABA_B_, *opioid µ and k, and cannabinoid CB1 receptors*.
- *GABAergic presynaptic terminals are most potently regulated by these receptors*.
- *Changes in the strength of these modulatory pathways may contribute to increased PB excitability and, consequently, chronic pain*.

## 1. Introduction

The parabrachial nucleus (PB) subserves sensory, homeostatic and aversive functions (Palmiter, 2018; Chiang et al., 2019, 2020), and is a critical hub for nociception (Gauriau and Bernard, 2002; Han et al., 2015). The role of PB in pain processing is highlighted by its reciprocal connections with brain regions associated with both the sensory and affective aspects of pain (Fulwiler and Saper, 1984; Jasmin et al., 1997; Chen et al., 2017) as well as with regions of descending modulatory control of nociception (Roeder et al., 2016; Chen et al., 2017; Chen and Heinricher, 2019).

In addition to normal nociception, PB also contributes to pathological pain conditions. Using a combination of rat and mouse models, we have shown that chronic pain is causally related to amplification of PB responses, and to reduced inhibition of PB neurons by the central amygdala (Uddin et al., 2018; Raver et al., 2020). These findings suggest that mechanisms that regulate the efficacy of synaptic transmission within PB, as well as the intrinsic excitability of PB neurons, significantly contribute to normal and dysregulated nociception. How this regulation occurs, however, has not yet been determined.

Several neurotransmitter receptors commonly associated with nociception as well as modulation of synaptic transmission are expressed in PB, including those activated by GABA_B_, µ- and k opioid (MOP/KOP), and cannabinoid type-1 (CB1) receptors. While the presence of these signaling pathways is known, their functional impact on PB neurons is not well established. In particular, little is known about how these receptors are distributed between inhibitory and excitatory afferents or whether their effects are biased towards pre- or postsynaptic regulation. Our ability to understand how PB contributes to chronic pain requires understanding how these neuromodulators affect synaptic release and neuronal excitability under normal conditions. This is the goal of the present study.

## 2. Results

### 2.1 GABAB_B_ receptor activation inhibits synaptic release

We have previously shown that GABAergic inputs to the parabrachial nucleus regulate the expression of pain behaviors. Here, we investigated the role of GABA_B_ receptors by recording pharmacologically isolated mIPSCs and mEPSCs before and after bath application of increasing concentrations of baclofen (0.1 to 300 µM). As shown in the representative mIPSC recordings in Figure 1A, 100 µM baclofen reduced the frequency of synaptic events. We obtained a dose-response profile by normalizing the median mIPSC frequency at each concentration of baclofen to the baseline value for each neuron. Only neurons that had a significant response to the agonist —determined by Kruskal-Wallis test with p < 0.05—were included (mIPSCs: n = 8 out of 9, mEPSCs: n = 5 out of 5 neurons).The stochastic nature of instantaneous event frequencies (CV: 270 ± 80 %, n = 11) and amplitudes (CV: 60 ± 20 %, n = 11) in each neuron is reflected in the normalized dose response group data variability at each concentration. Despite the inherent variability of these parameters, this analysis (Fig. 1B) revealed an IC50 of 1 µM (95% CI: 0.3 to 4 µM) and a maximum inhibition to 23% (95% CI: 11 to 34%) of baseline activity.

**Figure 1:**
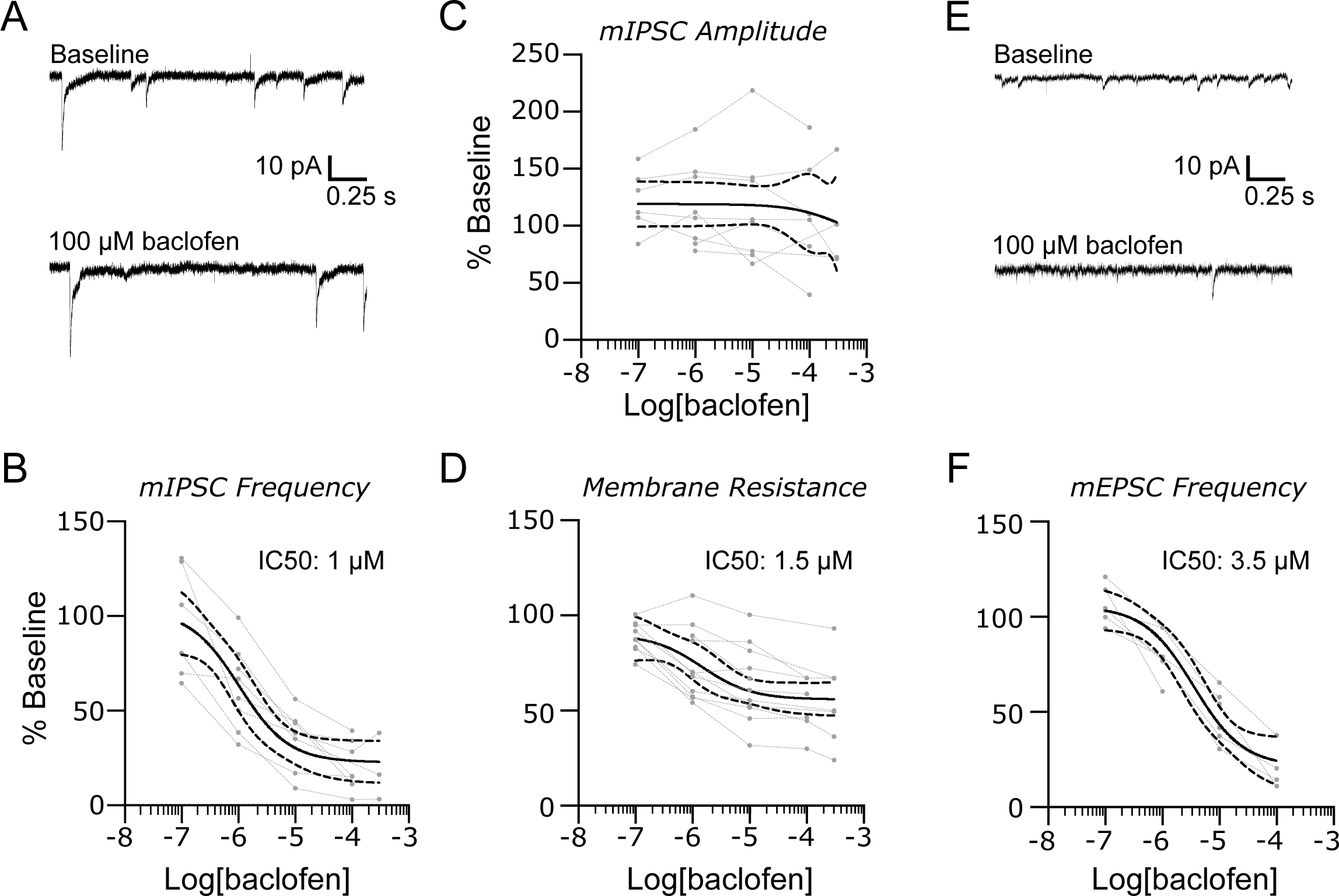
GABAB receptors suppress synaptic input and membrane resistance in the lateral parabrachial nucleus. (A) Representative recordings of mIPSCs in baseline conditions and in the presence of 100 mM baclofen. The suppression of mIPSC frequency was dose dependent (B). The amplitudes of mIPSC were not similarly affected (C) despite a dose-dependent reduction in PB neurons (D). Baclofen also reduced the frequency of mEPSCs. Representative recordings are shown in (E) with group data for mEPSC frequency in (F). Group data are fit with a log(dose) vs response curve ± 95% CIs.

The amplitudes of mIPSCs was significantly altered by baclofen in 6 out of 9 neurons (defined as above for each cell with a Krusal-Wallis (K-W) test with p <0.05), but the direction of change was not consistent across neurons. Median amplitudes recorded from some neurons decreased as a function of baclofen concentration, while smaller amplitude events were suppressed in other neurons resulting in an overall increase in the median amplitude. As a result, when examined as a population, there was no consistent effect of baclofen on mIPSC amplitude (Fig. 1C; K-W test, K-W test, H(4) = 2, P = 0.73), despite a dose dependent decrease in membrane resistance (Fig. 1D; K-W test, H(4) = 17, P = 0.0019: IC50 = 1.5 µM; 95% CI: 0.9 to 23 µM), with a maximum decrease to 56% (95% CI: 44 to 65%) at the highest baclofen concentration of 300 µM.

Baclofen had a similar inhibitory impact on mEPSC frequency in all 7 neurons recorded (Fig. 1E). Analysis of normalized mEPSC frequency group data (Fig. 1F) revealed a significant effect of baclofen (K-W test, K-W test, H(4) = 17.38, P = 0.006) with an IC50 of 3.5 µM (95% CI: 1 to 10 µM) and maximum suppression to 22% (95% CI: 3 to 36%) of baseline. We did not analyze mEPSC amplitude in these recordings because the cesium-based pipette solution blocks potassium channels and alters the membrane resistance.

Together, these data demonstrate that GABA_B_ receptors have both pre- and postsynaptic effects in PB by inhibiting GABAergic and glutamatergic transmission and activating a postsynaptic conductance.

### 2.2 µ and k opioid receptors differentially affect mEPSCs and mIPSCs

Mu opioid peptide (MOP) receptors are highly expressed in PB, suggesting that endogenous and exogenous agonists may also regulate synaptic activity in this nociceptive hub. We tested this prediction directly by recording mIPSCs and mEPSCs in the presence of increasing concentrations of the selective MOP agonist, DAMGO (1 nM to 1 µM). Representative mIPSC recordings in Figure 2A demonstrate a suppression of synaptic release probability by MOP receptor activation. Normalization of the median mIPSC frequencies following DAMGO application to baseline values revealed a significant. dose-dependent effect with an IC50 of 10 nM (95% CIs: 2.1 to 43 nM, 11 out of 13 neurons responding; K-W test, K-W test, H(3) = 16.47, P = 0.0009) and maximum inhibition to 46% of baseline values (95% CIs: 34 to 56 %, Fig. 2B). There was no effect of DAMGO on mIPSC amplitudes when examined as a population (Fig. 2C, K-W test, H(3) = 6.3, P = 0.1), despite a significant effect in 6 out of 13 neurons. However, DAMGO had a dose-dependent effect on postsynaptic membrane resistance with an IC50 of 110 nM (Fig. 2D, 95% CIs: 4nM to 9 µM, K-W test, H(4) = 9.6, P = 0.048). The membrane resistance was maximally reduced to 58% of the baseline value (upper 95% CI: 68%) at 3 µM DAMGO.

**Figure 2:**
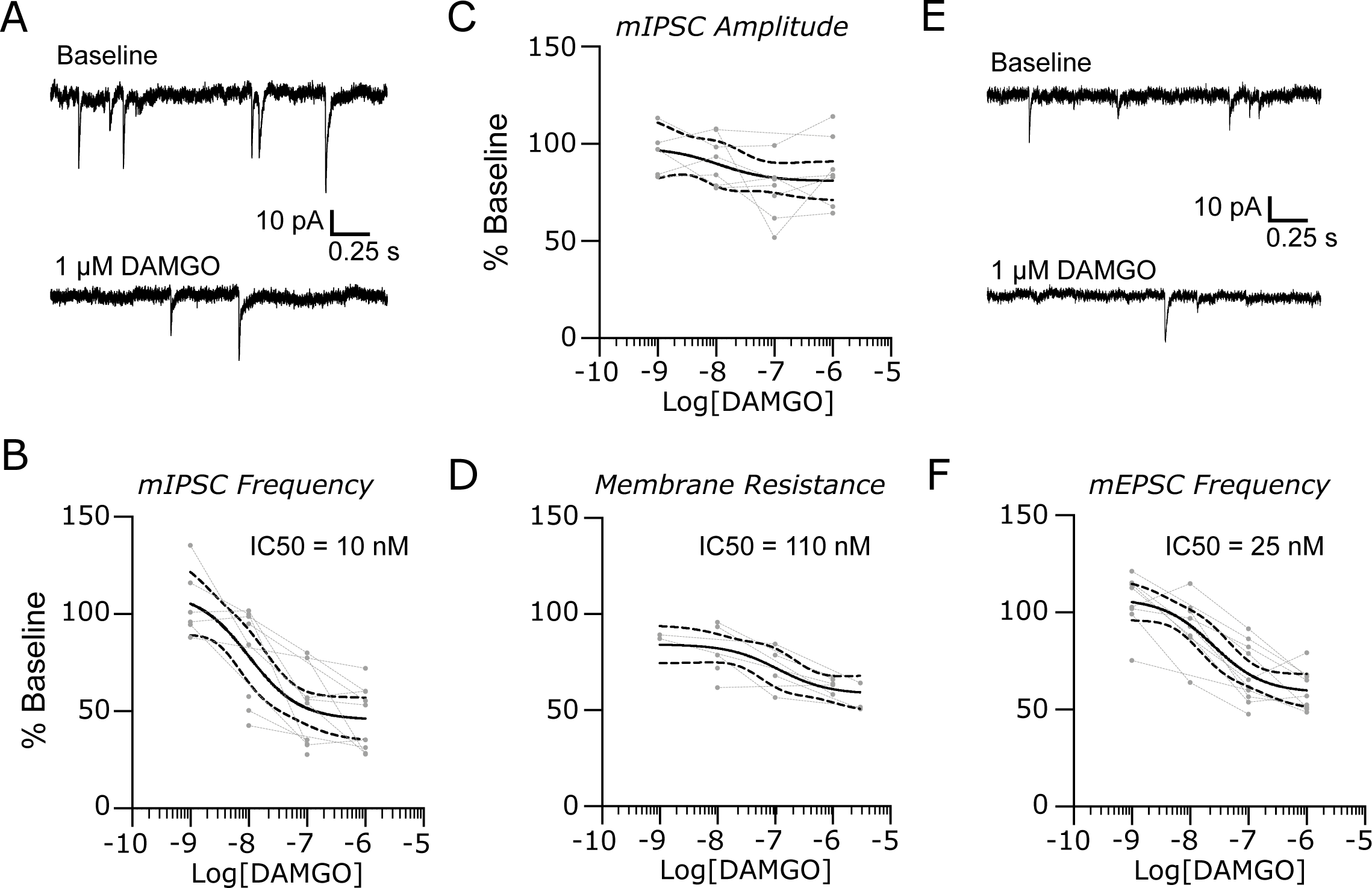
MOP receptors suppress synaptic activity and input resistance in the lateral parabrachial nucleus. (A) Representative recordings of mIPSCs in baseline conditions and in the presence of 1 µM DAMGO. The suppression of mIPSC frequency was dose dependent (B), but the amplitudes (C) were not affected. However, DAMGO caused a significant dose dependent reduction in membrane resistance in the PB neurons (D). Similar results were observed for mEPSCs with representative recordings in (D) and group data for frequency in (F). Group data are fit with a log(dose) vs response curve ± 95% CIs.

DAMGO had a similar inhibitory effect on the release probability of mEPSCs (Figs. 2E & F, K-W test, H(3) = 21.8, P < 0.0001) with an IC50 of 25 nM (95% CI: 7 to 86 nM, 13 out of 20 neurons responding).

Thus, like GABA_B_ receptors, MOP receptor activation has both pre- and postsynaptic affects. However, postsynaptic MOP receptors have a higher IC50 for DAMGO relative to presynaptic, and inhibitory inputs tend to be suppressed at lower concentrations relative to excitatory synapses.

Kappa opioid peptide receptors (KOP) are frequently associated with the aversive aspects of opioid signaling, despite their analgesic capacity (Cahill et al., 2014). Thus, their expression in PB may contribute to the negative aspects of nociception and to chronic pain conditions in particular. In contrast to the relatively consistent effects observed with GABA_B_ and MOP activation, the selective agonist, U-69593, differentially affected synaptic release probability at inhibitory and excitatory synapses.

The effect of U-69593 on mIPSCs was similar to that observed with baclofen and DAMGO. Representative recordings in Figure 3A demonstrate the suppression of mIPSC frequency following application of U-69593. Analysis of group data (Fig. 3B) revealed a dose dependent effect, with an IC50 of 16 nM (95% CIs: 1 to 200 nM, 7 out of 9 neurons responding, K-W test, H(4) = 11.5, P = 0.021). There was no effect of U69593 on either mIPSC amplitude (Fig. 3C, K-W test, H(4) = 2.2, P = 0.75) or membrane resistance (Fig. 3D, K-W test, H(4) = 1.4, P = 0.84).

**Figure 3:**
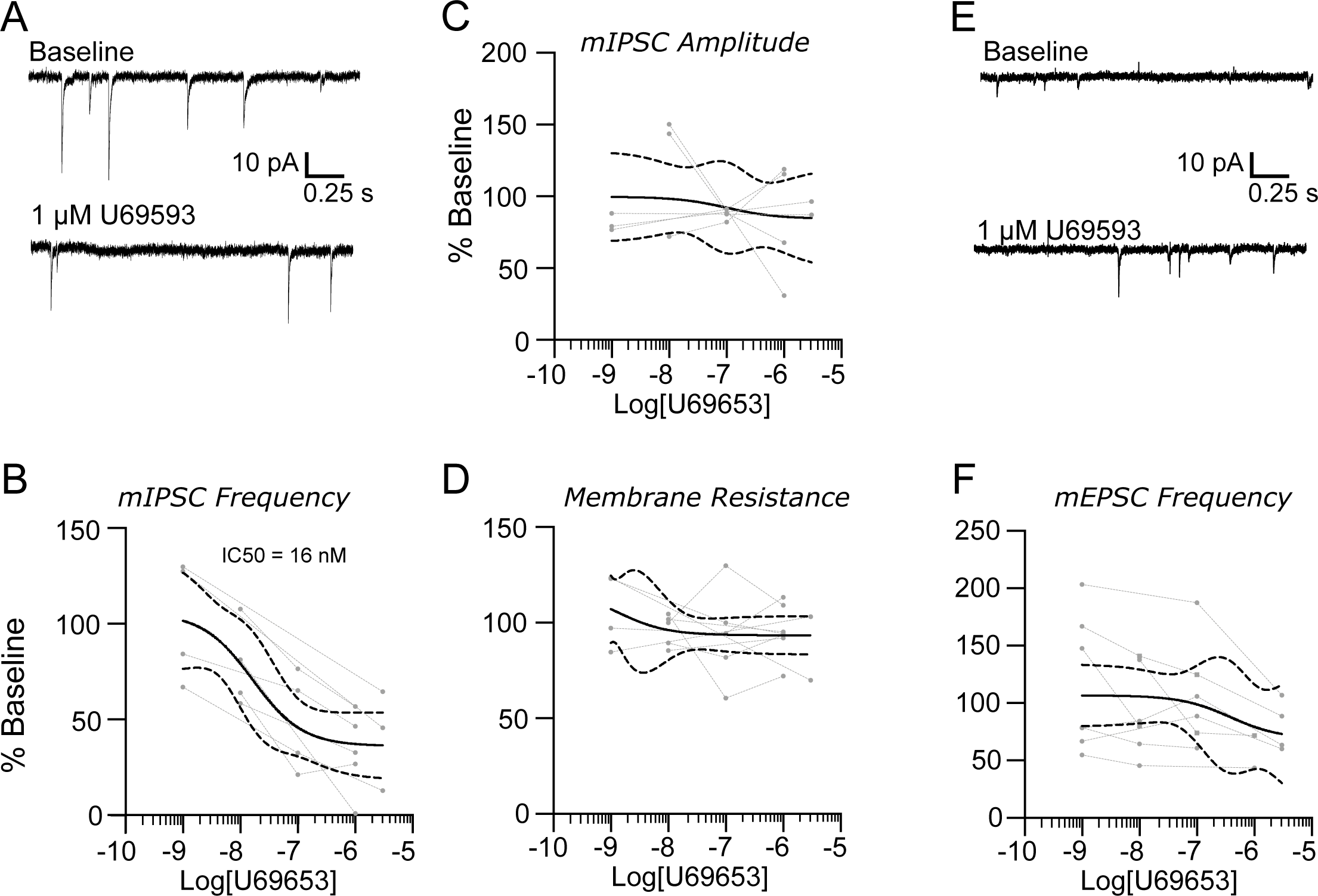
KOP receptors suppress inhibitory, but not excitatory, network activity in the lateral parabrachial nucleus. (A) Representative recordings of mIPSCs in baseline conditions (top) and in the presence of 1 µM U69593. The suppression of mIPSC frequency was dose dependent (B), but the amplitudes (C) were not affected. There was also no effect of KOP receptor activation on neuronal membrane resistance (D). Representative recordings of mEPSCs (E) and group data (F) show no consistent effect of U69593 on the frequency of excitatory events. Group data are fit with a log(dose) vs response curve ± 95% CIs.

In contrast, KOP activation had highly variable effects on mEPSC frequencies. We observed a reduction in mEPSC frequency in 5 out of 14 neurons, no significant change in 4 neurons, and the remaining 5 neurons showed a significant increase in frequency in the presence of U-69593. Sample recordings for a neuron where mEPSC frequency increased in response to this agonist are shown in Figure 3E. Group data are shown in Figure 3F, where neurons with a significant decrease in mEPSC frequency are depicted with solid lines and those with a significant increase are indicated with dashed lines. Both types of responses lacked a clear dose-dependent relationship, and, as a population, did not result in a meaningful IC50 value (K-W test, H(4) = 3.71, P = 0.45).

### 2.3 Endocannabinoids modulate excitatory and inhibitory synaptic release

Cannabinoid type 1 (CB1) receptors are widely expressed in the brain, including nuclei involved in nociception such as PB (Herkenham et al., 1991). We investigated the role of cannabinoid signaling by recording pharmacologically isolated mIPSCs and mEPSCs before and after bath applying increasing concentrations of WIN 55,212-2 (WIN: 0.1 to 50 µM), a CB1 receptor agonist. As observed in the representative recordings of mIPSCs in Figure 4A, 50 µM WIN decreased the frequency of synaptic events (K-W test, H(5) = 19.7, P = 0.0015). Dose-response analysis for the 13 of 16 neurons affected yielded an IC50 of 400 nM (Fig. 4B, 95% CI: 60 nM to 1.4 µM). As a population, median mIPSC amplitudes were not consistently altered by WIN (Fig. 4C, K-W test, H(5) = 4.63, P = 0.46), and we did not observe a consistent change in membrane resistance (Fig. 4D, K-W test, H(4) = 0.77, P = 0.94).

**Figure 4:**
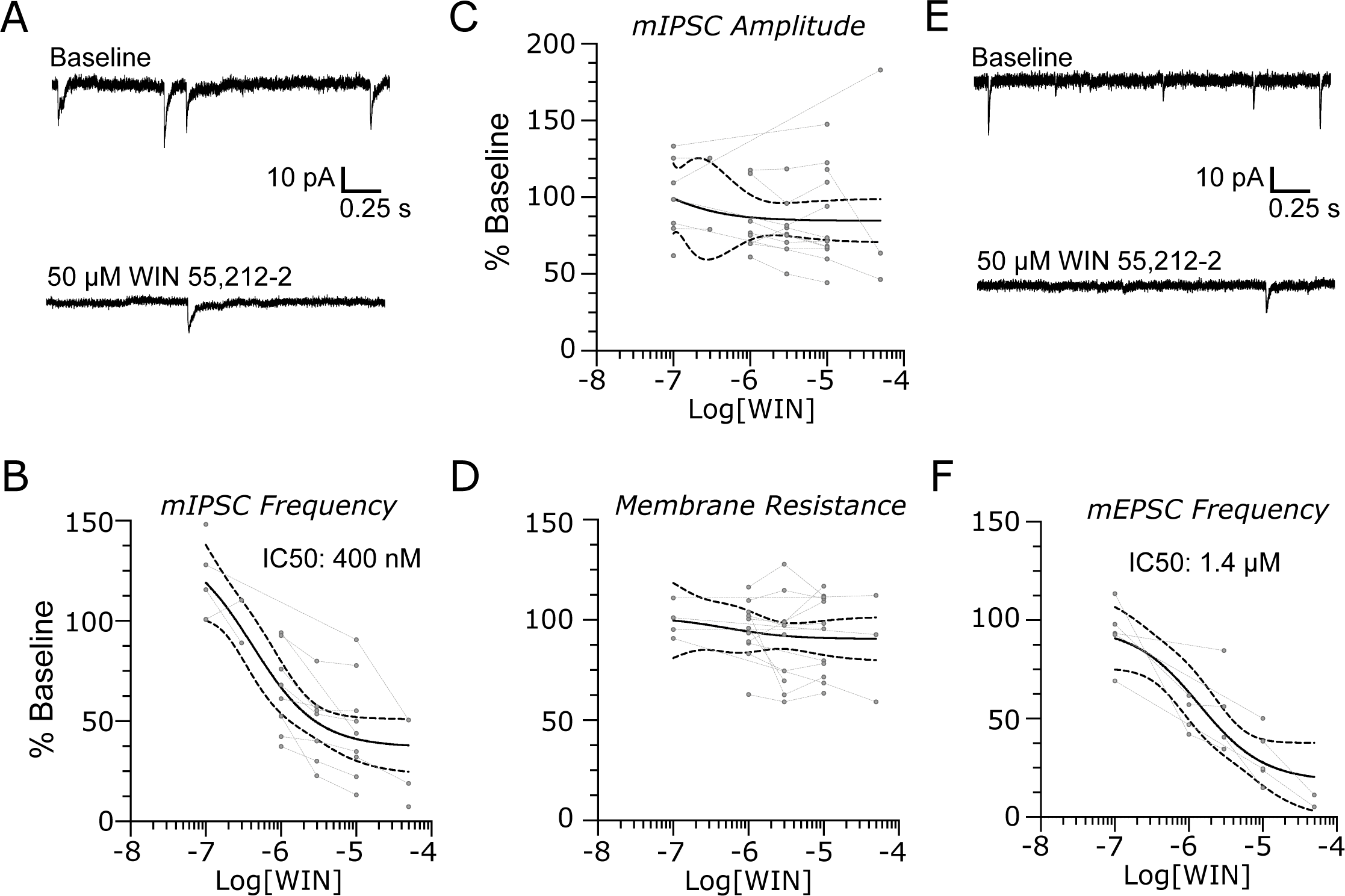
Endocannabinoid receptors modulate synaptic activity in the lateral parabrachial nucleus. (A) Representative recordings of mIPSC at baseline (top) and in the presence of 50 μM WIN 55,212-2. The CB1R agonist caused a significant dose dependent decrease in mIPSC frequency (C) without consistently affecting the event amplitudes (D) or postsynaptic membrane resistance (D) Representative recordings of mEPSC at baseline and in the presence of 50 µM WIN 55,212-2 show a reduction in excitatory event frequency (E) with a dose-dependence (F) Group data are fit with a log(dose) vs response curve ± 95% CIs.

There was a similar inhibitory effect of WIN on mEPSC frequency in 8 out of 9 PB neurons. Representative traces are shown in Figure 4E. Analysis of normalized mEPSC frequency group data for these 8 neurons yielded an IC50 of 1.4 µM (Fig. 4F, 95% CI: 0.1 to 10 µM, K-W test, H(4) = 15.3, P = 0.0042).

Together, these data support a role for CB1 receptors in regulation of synaptic release, without a direct postsynaptic effect.

### 2.4 Endocannabinoid receptors are tonically active in the lateral parabrachial nucleus

Tonic endocannabinoid activity is frequently observed at CB1 receptors, and changes in the level of this tonic activity have been reported in models of chronic pain (Dogrul et al., 2002).Using the specific CB1 receptor antagonist AM251, we tested whether similar tonic activity is present in PB. At inhibitory synapses, applying increasing concentrations of AM251 (0.1 to 10 nM) led to a dose-dependent increase in events in 13 out of 14 neurons (Fig. 5A) with an EC50 of 1.4 nM (95% CIs: 0.04 to 70 nM, K-W test, H(3) = 9, P = 0.03).

**Figure 5:**
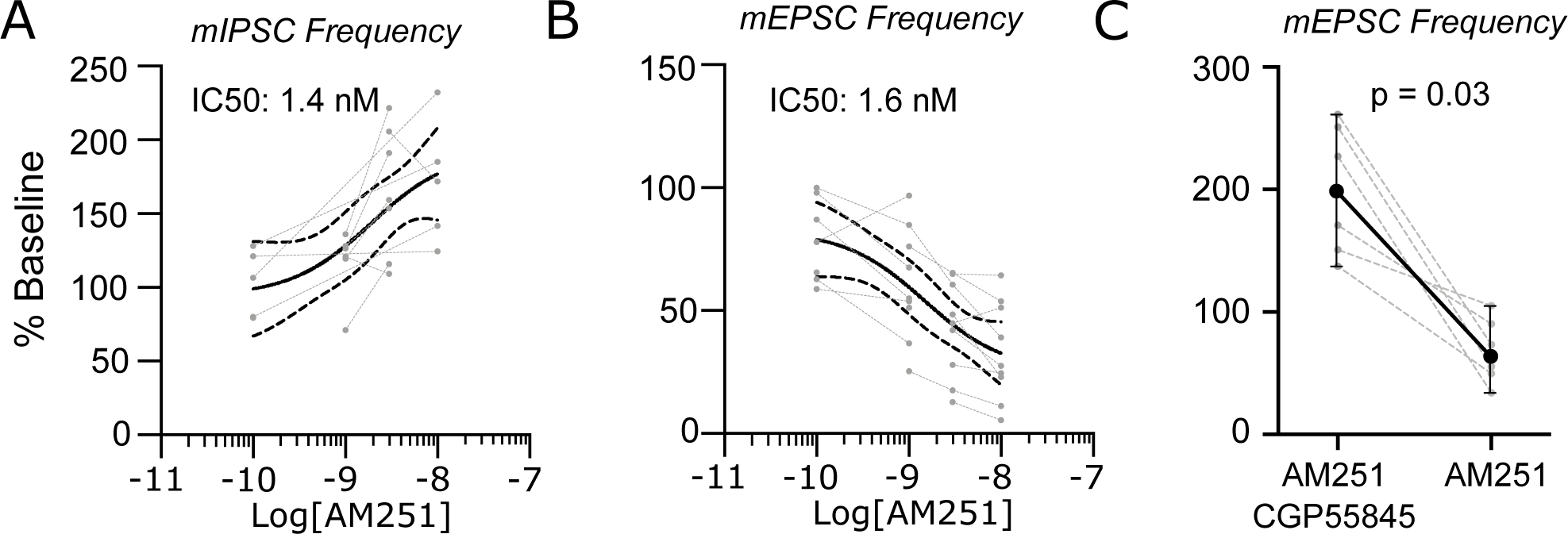
Endocannabinoid receptors are tonically active in the lateral parabrachial nucleus. (A) AM251 caused a significant dose dependent increase in mIPSC frequency but had the opposite effect on mEPSC frequency (B). This effect on mEPSC frequency was reversed by the co-application 1µM CGP55845, a GABAB receptor antagonist (C). Dose response data are fit with a log(dose) vs response curve ± 95% CIs.

In contrast to the effect on mIPSCs, and surprisingly, AM251 led to a dose-dependent *decrease* of mEPSC frequencies, with a similar IC50 of 1.6 nM (95% CIs: 0.2 to 30 nM, Fig. 5B, K-W test, H(3) = 12.9, P = 0.0048). Because this reduction occurs during an AM251-driven increase in mIPSC release, we hypothesized that activation of GABA_B_ receptors on glutamatergic synapses may drive this counterintuitive result. We tested this hypothesis by blocking GABA_B_ receptors with 1 µM CGP55845, a selective GABA_B_ receptor antagonist, before applying 3 nM AM251. This resulted in an increase in mEPSC frequency, similar to that observed for GABAergic synapses (Fig. 5C). The net inhibitory effect of AM251 on mEPSC frequency was restored after washout of CGP55845 (Paired Wilcoxon test, p = 0.03, n = 6 neurons). This finding suggests that the efficacy of tonic CB1 signaling at glutamatergic synapses in PB is regulated by GABA_B_ receptors.

## 3. Discussion

We tested the hypothesis that signaling pathways associated with nociceptive processing would modulate synaptic activity in the parabrachial nucleus. We focused on the GABAergic system, which we have shown to be dysregulated in PB in a model of chronic pain (Uddin et al., 2018; Raver et al., 2020), and on two pathways closely associated with antinociception – the endogenous opioid and cannabinoid systems. Our results indicate that graded activation of GABA_B_, µ- and *k*-opioid and CB_1_ receptors suppress synaptic release in PB. We find that GABAergic and glutamatergic synapses in this nucleus are under tonic CB1R-mediated suppression, but that the impact of CB1R signaling at excitatory synapses is regulated by GABA_B_ receptors. Additionally, we find that IC50s for inhibitory inputs tended to be lower than those for excitatory suggesting these pathways are biased towards regulating inhibitory synapses. These differences offer insights into the functional significance these pathways have in the regulation of PB excitability, and in nociception.

### 3.1 GABA_B_ receptors diminish pre- and postsynaptic excitability

We show that the GABA_B_ receptors regulate presynaptic release of both glutamate and GABA from synapses with PB neurons. The presynaptic receptors had similar affinities to baclofen. Because synaptically released GABA typically has to diffuse for longer distances to affect glutamatergic synapses—compared to the shorter distances to affect GABA_B_ auto-receptors—it is likely that GABA has a more potent effect on inhibitory than on excitatory synapses in PB. In line with prior reports (Christie and North, 1988) we demonstrate that activation of these receptors also reduced the input resistance of PB neurons, consistent with a postsynaptic effect. Thus, GABA_B_ mediated signaling likely suppresses nociceptive transmission in PB in normal conditions. In pathological pain, however, this role may be compromised.

We have recently shown that PB receives dense GABAergic innervation from the central nucleus of the amygdala (CeLC, the “nociceptive amygdala”), and that, in an animal model of chronic pain, this inhibitory pathway is suppressed (Raver et al., 2020). We also demonstrated that this suppression is causally related to chronic pain (Raver et al., 2020). The suppressed release of GABA may lead to the amplified activity of PB neurons, seen in chronic pain conditions (Uddin et al., 2018; Raver et al., 2020) through at least two mechanisms. Reduced GABA release may lead to reduced activation of postsynaptic GABA_B_ receptors, resulting in dis-inhibition of PB neurons and amplifying responses to nociceptive inputs (Uddin et al., 2018; Raver et al., 2020).

Reduced activation of GABA_B_ may also lead to more profound postsynaptic changes. For example, in the spinal cord, GABA_B_ receptors are essential for modulating after-discharges (Russo et al., 1998). After-discharges are prolonged neuronal responses that outlast a sensory stimulus (Woolf and King, 1987; Herrero et al., 2000). The durations of after-discharges and the proportion of neurons that express them, is dramatically increased in chronic pain (Palecek et al., 1992; Laird and Bennett, 1993). We reported that, in PB of both rats and mice with chronic pain, the incidence and duration of after-discharges is markedly increased(Uddin et al., 2018; Raver et al., 2020). After-discharges may be causally related to the expression of chronic pain (Laird and Bennett, 1993; Asada et al., 1996). For example, we have demonstrated that the incidence and duration of after-discharges in spinal neurons increases significantly in animals with chronic, neuropathic pain, and that suppressing after-discharges significantly lessens hyperalgesia in experimental animals (Okubo et al., 2013). Thus, the suppressed GABA release in PB during chronic pain (Charlie’s paper) likely results in reduced activation of postsynaptic GABA_B_ receptors, resulting in generation of after-discharges and chronic pain.

### 3.2 MOP and KOP have mixed effects on PB excitability

The high expression levels of µ-opioid peptide (MOP) receptors in PB (Mansour et al., 1994), including in neurons that project to the amygdala (Chamberlin et al., 1999), suggests that these receptors are key modulators of neuronal activity in PB and contribute to the affective aspect of nociception. We find that the selective MOP agonist, DAMGO, suppresses presynaptic release of both GABA and glutamate in a dose dependent manner, indicating a reduced probability of release at these synapses. DAMGO affected inhibitory synapses at lower concentrations compared to excitatory ones (IC50s: 9 vs 50 nM), suggesting that low levels of agonist activity may facilitate nociceptive transmission in PB. We also find that DAMGO activates a postsynaptic conductance with an IC50 of 110 nM, similar to values reported for DAGOL (Christie and North, 1988). Although not directly tested here, this conductance was determined to result in an inwardly rectifying potassium current (Christie and North, 1988). Together, these results suggest that, as MOP receptor activity increases, presynaptic effects precede postsynaptic ones, and that low concentrations of agonist may increase nociceptive transmission by preferentially suppressing inhibitory transmission.

A deeper understanding of how this differential regulation affects nociception is important given the complexity of responses of MOP receptors to chronic disease conditions. For example, MOP receptor activity in PB is reduced following chronic opioid exposure (Sim et al., 1996; Sim-Selley et al., 2000), while receptor expression levels are increased following chronic constriction injury of the sciatic nerve (Llorca-Torralba et al., 2020). Yet, both opioid withdrawal and nerve injury are associated with chronic pain.

Similar to MOP agonists, KOP agonists are widely expressed throughout the brain, including PB (Mansour et al., 1994), where they play a critical role in mediating the aversive aspects of nociception (Chiang et al., 2020). The association between KOP signaling and aversive behavioral effects, such as dysphoria and respiratory depression (Pfeiffer et al., 1986; Roth et al., 2002; Wadenberg, 2003; Land et al., 2008; Tejeda et al., 2013), have limited the therapeutic use of exogenous agonists, despite their ability to mediate analgesia with less risk of abuse (Wang et al., 2010). Our findings support an aversive role for KOP signaling in nociception. We find that KOP agonists preferentially suppress synaptic release at GABAergic synapses and leave glutamatergic signaling relatively intact. This bias towards suppression of inhibition may facilitate propagation of nociceptive signals to higher brain centers such as the amygdala (Chiang et al., 2020). The reported lack of a postsynaptic effect of KOP receptors on PB neuronal excitability (Christie and North, 1988), which we also observed here, further supports a faciliatory role for PB KOPs in nociceptive signaling. In a model of chronic pain, no changes were reported in KOP expression levels, despite significant increases in MOP expression (Llorca-Torralba et al., 2020). Whether there are changes in the strength of this pathway during chronic pain remain to be determined.

### 3.3 Cannabinoid signaling selectively regulates presynaptic activity in PB

Cannabinoid type 1 receptors (CB1R) are G-protein coupled receptors (GPCRs) widely expressed in the brain, including PB (Herkenham et al., 1991). Activation of this pathway is antinociceptive, consistent with their ability to suppress presynaptic activity via retrograde signaling from the postsynaptic neuron (Alger, 2002; Manzanares et al., 2006; Woodhams et al., 2017; Vuckovic et al., 2018). In PB we find that activation of CB1R with the agonist WIN 55,212-2 reduces probability of release at both glutamatergic and GABAergic synapses, with no significant impact on the intrinsic excitability of PB neurons. As with the other agonists investigated here, regulation of synaptic activity by CB1R is biased towards suppression of inhibition, suggesting that expression may be higher in inhibitory synapses, as reported in other brain regions (Kano et al., 2009).

Tonic CB1R activity mediates antinociception in the spinal cord, and its loss results in lowered pain thresholds (Dogrul et al., 2002). We find that CB1R in PB are tonically active, as application of the CB1R antagonist AM251 produced a dose-dependent increase in synaptic release at GABAergic synapses. In contrast, AM251 had the opposite effect on excitatory synapses and reduced the frequency of mEPSCs, an effect that was reversed by blocking GABA_B_ receptors. Thus, CB1 receptors in PB are tonically active at inhibitory and excitatory synapses, but the net impact on the latter is modulated by GABA. This suggests that, under normal circumstances, GABA_B_ signaling prevents shifts in tonic CB1R activity from increasing excitation in PB. Future investigations will examine if the reduction in GABAergic signaling observed in chronic pain removes this brake and allows shifts tonic CB1 activity to increase excitatory transmission in a pronociceptive manner.

Together, our results provide important insights into neuromodulatory control of synaptic transmission and excitability within PB and provide a foundation for future studies on how changes in these pathways may contribute to chronic pain.

## 4. Materials and Methods

### 4.1 Animals

All animal procedures were reviewed and approved by the University of Maryland Institutional Animal Care and Use Committee and adhered to the National Institutes of Health guide for the care and use of laboratory animals and ARRIVE guidelines. We used male and female adult (∼7 to 13 weeks) C57Bl6/J (n = 64, Jackson Laboratory) mice from our in-house colony. For experiments testing the effects of DAMGO we used C57Bl6/J TRPV1-ChR2 (n = 17) generated by crossing Ai32(ChR2/EYFP) with TRPV1^Cre^ mice. These mice were generated as part of an independent study. Because the frequency of synaptic events in these animals was indistinguishable from that in C57Bl6/J mice (p≥0.34, Mann-Whitney U) we combined data from these strains. Similarly, because there the frequency of synaptic events in males and females was indistinguishable (p=0.68, Mann-Whitney U) we combined data from both sexes.

### 4.2 Slice preparation

Animals were deeply anesthetized with ketamine (180 mg/kg) and xylazine (20 mg/kg), and the brains were rapidly removed following decapitation. Sagittal slices through PB, 300 µm thick, were cut in ice-cold cutting artificial cerebral spinal fluid (ACSF) using a Leica VT1200s vibratome (Leica Biosystems, Buffalo Grove, IL) and transferred to warm (32–34°C) recovery ACSF for 10–15 min. The slices were then transferred to normal ACSF at room temperature for at least 45 min before starting experiments. All solutions were continuously bubbled with a mixture of 95% oxygen and 5% CO_2_.

### 4.3 Solutions and drugs

ACSF compositions were based on the methods of Ting et al (Ting et al., 2014) and consisted of (in mM); cutting ACSF: 92 NMDG, 30 NAHCO_3_, 20 HEPES, 25 glucose, 5 Na-ascorbate, 2 thiourea, 1.25 NaH_2_PO_4_, 2.5 KCl, 3 Na-pyruvate, 0.5 CaCl_2_ and 10 MgSO_4_; normal ACSF: 119 NaCl, 2.5 KCl, 1.25 NaH_2_PO_4_, 24 NaHCO_3_, 12.5 glucose, 2 CaCl_2_ and 2 mM MgSO_4_. The pH and osmolarities of each were adjusted to 7.35–7.45 and 300–310 mOsm, respectively. Solutions were continuously saturated with carbogen (95% O_2_, 5% CO_2_) throughout use. For experiments targeting excitatory synaptic currents we used a pipette solution consisting of (in mM): 130 Cs-Methanesulfonate, 10 HEPES, 0.5 EGTA, 1 MgCl_2_, 2.5 Mg-ATP and 0.2 GTP-Tris. For targeting inhibitory and postsynaptic currents, we used a pipette solution consisting of (in mM): 70 K-Gluconate, 60 KCl, 10 HEPES, 1 MgCl_2_, 0.5 EGTA, 2.5 Mg-ATP, 0.2 GTP-Tris. Both pipette solutions were adjusted to a pH of 7.3 and 285 mOsm. Information on receptor agonists and antagonists are provided in Table 1.

**Table 1:** Receptor agonists and antagonists used for dose-response analysis.

| Drug | Solvent | Supplier | Catalog Number | Concentration |
| --- | --- | --- | --- | --- |
| Tetrodotoxin citrate | ddH<sub>2</sub>O | abcam | ab120055 | 0.5 to 1 µM |
| CNQX (6-cyano-7-nitroquinoxaline-2,3-dione) | ddH<sub>2</sub>O | Sigma-Aldrich | C239 | 20 µM |
| APV (DL-2-Amino-5-phosphonopentanoic acid) | ddH<sub>2</sub>O | Sigma-Aldrich | A5282 | 50 µM |
| Gabazine | ddH<sub>2</sub>O |  |  | 10 µM |
| Baclofen | 0.1 N NaOH | Tocris Bioscience | 0417 | 0.1 – 300 µM |
| DAMGO | ddH<sub>2</sub>O | abcam | ab120674 | 1 nM – 3 µM |
| U-69593 | 50% EtOH | abcam | ab141703 | 1 nM – 3 µM |
| WIN-55212-2 | DMSO | Axxora | BML-CR105 | 1 – 10 µM |
| AM251 | DMSO | Sigma-Aldrich | A6226 | 0.1 – 10 nM |
| CGP55845 | DMSO | Tocris Bioscience | 1248 | 1 µM |

### 4.4 Electrophysiology

Whole cell patch-clamp recordings were obtained from neurons in lateral PB with a Multiclamp 700B amplifier (Molecular Devices) low-pass filtered at 1.8 kHz with a four-pole Bessel filter, and digitized with Digidata 1550B (Molecular Devices). Lateral PB is easily identified in these slices by its proximity to the superior cerebral peduncle. Recording locations were identified visually at low magnification before and after each recording for all neurons and verified with biocytin immunohistochemistry. The impedance of patch electrodes was 4–8 MΩ. Once a GΩ seal was obtained, holding potential was set to −65 mV and was maintained for the duration of the experiment. All recordings were obtained at room temperature.

Miniature inhibitory presynaptic currents (mIPSCs) were recorded in the presence of 0.5 to 1 µM TTX and 10.0 µM gabazine, while miniature excitatory presynaptic currents (mEPSCs) were recorded in the presence of 0.5 to 1 µM TTX, 20 µM CNQX and 50 µM APV, respectively. We generated dose-response profiles by serial bath application of the respective agonist or antagonist, with a minimum 3 minute wash-in time per concentration, and collected only a single neuron per slice. All drugs and respective concentrations are provided in Table 1. Series resistance was monitored throughout the recordings with −5 mV hyperpolarizing pulses and we discarded recordings in which the resistance changed by more than 20% within a recording.

### 4.5 Biocytin Immunohistochemistry

Upon completion of recording, the pipette was carefully retracted from the neuron and the tissue slice transferred to 10% formalin at 4°C overnight. Slices were then washed 3 times for 10 minutes at room temperature in 1X PBS before being incubated overnight in a solution of 1:1000 Strepavidin – Cy3 conjugate, 3% fetal bovine serum and 0.3% Triton X-100 at 4°C. Slices were cover-slipped with an aqueous mounting media and visualized on a Leica SP8 confocal microscope to verify the recoding location within the lateral PB. We rejected data from 2 neurons which where determined to be outside this nucleus.

### 4.6 Data analysis and Statistics

Miniature inhibitory and excitatory postsynaptic currents were isolated offline using miniAnalysis (Synaptosoft). We used Clampfit (Molecular Devices) or the Neuromatic XOP for Igor (Wavemetrics) developed by Jason Rothman (Rothman and Silver, 2018) to calculate the membrane resistance based on the steady state current evoked by a −5 mV hyperpolarizing step. As the mIPSC and mEPSC frequencies and amplitudes were not normally distributed, we used the nonparametric Kruskal-Wallis test to check for effects of different agonist/antagonist concentrations within each neuron. We included only neurons which had a significant response (defined as p < 0.05) in the group data analysis. The fraction of neurons that responded significantly is indicated in the corresponding Results section. Group data for each drug and parameter were fit with a three parameter [inhibitor] vs response model in GraphPad Prism and, when the fit was successful, the IC50 with 95 % confidence intervals are reported.

## Author Contributions

**Nathan Cramer:** Conceptualization, Investigation, Formal analysis, Writing – Reviewing and Editing. **Gleice Cardoso:** Conceptualization, Investigation, Formal analysis, Writing – Original draft preparation, Funding acquisition. **Asaf Keller:** Conceptualization, Investigation, Writing – Reviewing and Editing, Funding acquisition, Resources

